# Sleep firing rate homeostasis is disrupted in mild parkinsonism

**DOI:** 10.1101/2025.04.18.649592

**Authors:** Bharadwaj Nandakumar, Ajay K. Verma, Ying Yu, Ethan Marshall, Adele DeNicola, Jing Wang, Colum D. MacKinnon, Michael J. Howell, Jerrold L. Vitek, Luke A. Johnson

**Affiliations:** Department of Neurology, University of Minnesota, Minneapolis, MN, USA

**Author notes:** Corresponding author Luke A. Johnson, PhD, Associate Professor, Department of Neurology, University of Minnesota, Lions Research Building 410, 2001 6^th^ ST SE, Minneapolis, MN, USA. These authors contributed equally (Bharadwaj Nandakumar, Ajay K. Verma).

## Abstract

Sleep-associated downscaling of neuronal firing, i.e., *firing rate homeostasis (FRH*), is essential for restorative sleep. Sleep dysfunction is common in Parkinson’s disease (PD), but FRH has not been investigated. Here, using a within-subject design in the nonhuman primate model of PD, we report that thalamocortical FRH is disrupted in parkinsonism. These findings can inform therapeutic approaches tailored towards normalizing FRH to reestablish restorative sleep in PD.

---

Sleep dysfunction is a common non-motor symptom in people with Parkinson’s disease (PD)^1^; however, effective treatment options to improve sleep remain elusive. A lack of understanding of the neuronal mechanisms underlying sleep dysfunction in PD is a major knowledge gap hindering the development of targeted therapies for enhancing sleep.

Firing rate homeostasis (FRH), characterized as the downscaling of neuronal firing (i.e., reduction in firing rate) from wake to sleep^2–4^ and a progressive reduction in neuronal firing from the time of sleep onset^5,6^, is an essential neural mechanism governing the restorative aspect of sleep^2^. This mechanism is associated with a sleep-related increase in the amplitude of cortical slow wave activity (SWA 0.1-4 Hz) and is crucial for maintaining diurnal energy stability at the cellular level^3,7^. A disruption in the FRH mechanism could compromise the restorative aspect of sleep and adversely impact behaviors dependent on sleep quality, such as learning, cognition, and memory^3,8^.

Sleep dysfunction in advanced PD is often manifested by attenuation in SWA during sleep and is associated with cognitive decline^9,10^, faster disease progression^11^, and excessive daytime sleepiness^12^. Our recent work suggests that sleep could also be disrupted in the early stage of parkinsonism^13^; however, the relationship between disrupted sleep and FRH in the early disease stage remains to be investigated. An improved understanding of the neuronal mechanisms underlying sleep dysfunction in the early disease stage could inform the development of targeted treatments for normalizing sleep quality in PD. To this end, we characterized sleep-related modulation in neuronal firing of the primary motor cortex (MC) and motor thalamus (MTh), key nodes of the basal ganglia thalamocortical network^14,15^ implicated in the manifestation of cardinal motor signs as well as sleep dysfunction in PD^16,17^. We hypothesized that disruption in SWA in mild parkinsonism would be associated with dysfunctional FRH, i.e., attenuation in wake-to-sleep downscaling of neuronal firing and impairment in progressive downscaling of neuronal firing during sleep.

An example spectrogram highlighting the disruption of cortical SWA during sleep in mild parkinsonism is shown in **Figure 1a**. We found that the SWA amplitude during sleep, an indicator of sleep quality, was markedly lower (WRS test, p=0.004) in mild parkinsonism compared to normal (**Figure 1b**). Next, we sought to characterize the effect of mild parkinsonism on the downscaling of neuronal firing of cortical and thalamic neurons from wake to sleep. An example highlighting wake-to-sleep firing rate changes in the cortical and thalamic neurons in normal and mildly parkinsonian states is shown in **Figure 1c**. The proportion of cortical neurons showing a reduction in firing rate between the wake and sleep states was reduced (Fischer’s test, p=0.003) in mild parkinsonism compared to normal (**Figure 1d**). Furthermore, wake-to-sleep modulation in firing rate, i.e., depth of modulation (DM), was lower (WRS test, p=0.005) in mild parkinsonism compared to normal (**Figure 1e**). The proportion of thalamic neurons reducing their firing rate from wake to sleep was not different (Fischer’s test, p=0.13) between normal and mildly parkinsonian states (**Figure 1f**). The DM in thalamus, however, was lower (WRS test, p<0.001) in mildly parkinsonian state compared to normal (**Figure 1g**).

**Figure 1.**
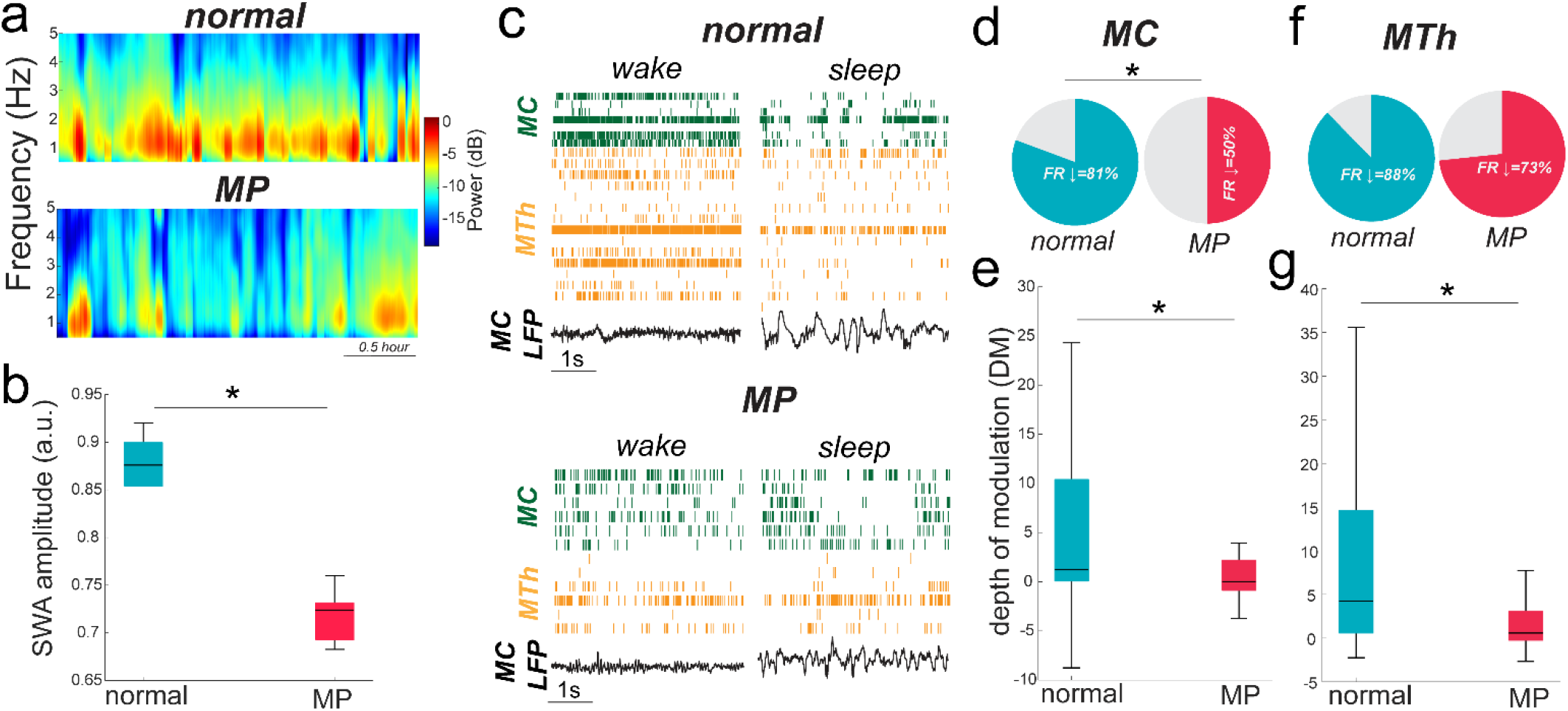
Example spectrograms showing a reduction in power of slow cortical oscillations during the first sleep cycle in mild parkinsonism (MP) compared to normal **(a)**. The distribution of SWA amplitude during sleep in normal (n=6) and MP (n=7) is shown in **(b)**. The amplitude of SWA during sleep was attenuated in MP compared to normal. Example recordings of single unit activity in MC and MTh and average MC LFP during wake and sleep in normal (top) and MP (bottom) reveal loss of firing rate reduction from wake to sleep (SWS) in MP **(c)**. The proportion of cortical neurons showing a reduction in firing rate (FR) from wake to SWS was attenuated in MP (36/72) compared to normal (50/62) **(d)**. The wake to SWS DM of cortical neurons was found to be attenuated in MP (n=72) compared to normal (n=62) **(e)**. The proportion of MTh neurons that reduced their FR from wake to SWS was not different between normal (57/65) and MP (22/30) **(f)**, however, wake to SWS DM of MTh neurons was attenuated in MP (n=30) compared to normal (n=65).

In addition to the downscaling of neuronal firing from wake to sleep, another aspect of FRH is the progressive downscaling of neuronal firing from the time of sleep onset. The effect of mild parkinsonism on FRH during sleep was evaluated by examining the firing rate-SWA and firing rate-time correlations. The proportion of neurons that exhibited a negative firing rate and SWA amplitude correlation during sleep was decreased in mild parkinsonism compared to normal for both MC (Fischer’s test, p<0.001) and MTh (Fischer’s test, p<0.001) neurons (**Figure 2a**). Additionally, the strength of firing rate-SWA correlation was found to be different (WRS test, p<0.001) between normal and mildly parkinsonian conditions for both cortical and thalamic neurons (**Figure 2b**). Similarly, the proportion of cortical (Fischer’s test, p=0.005) and thalamic (Fischer’s test, p=0.014) neurons that exhibited a negative correlation from the onset of sleep was decreased in mild parkinsonism compared to normal (**Figure 2c**). Moreover, the strength of firing rate-time (from the onset of sleep) correlation was different between normal and mild parkinsonian states for both MC (WRS test, p<0.001) and MTh (WRS test, p=0.04) neurons (**Figure 2d**).

**Figure 2.**
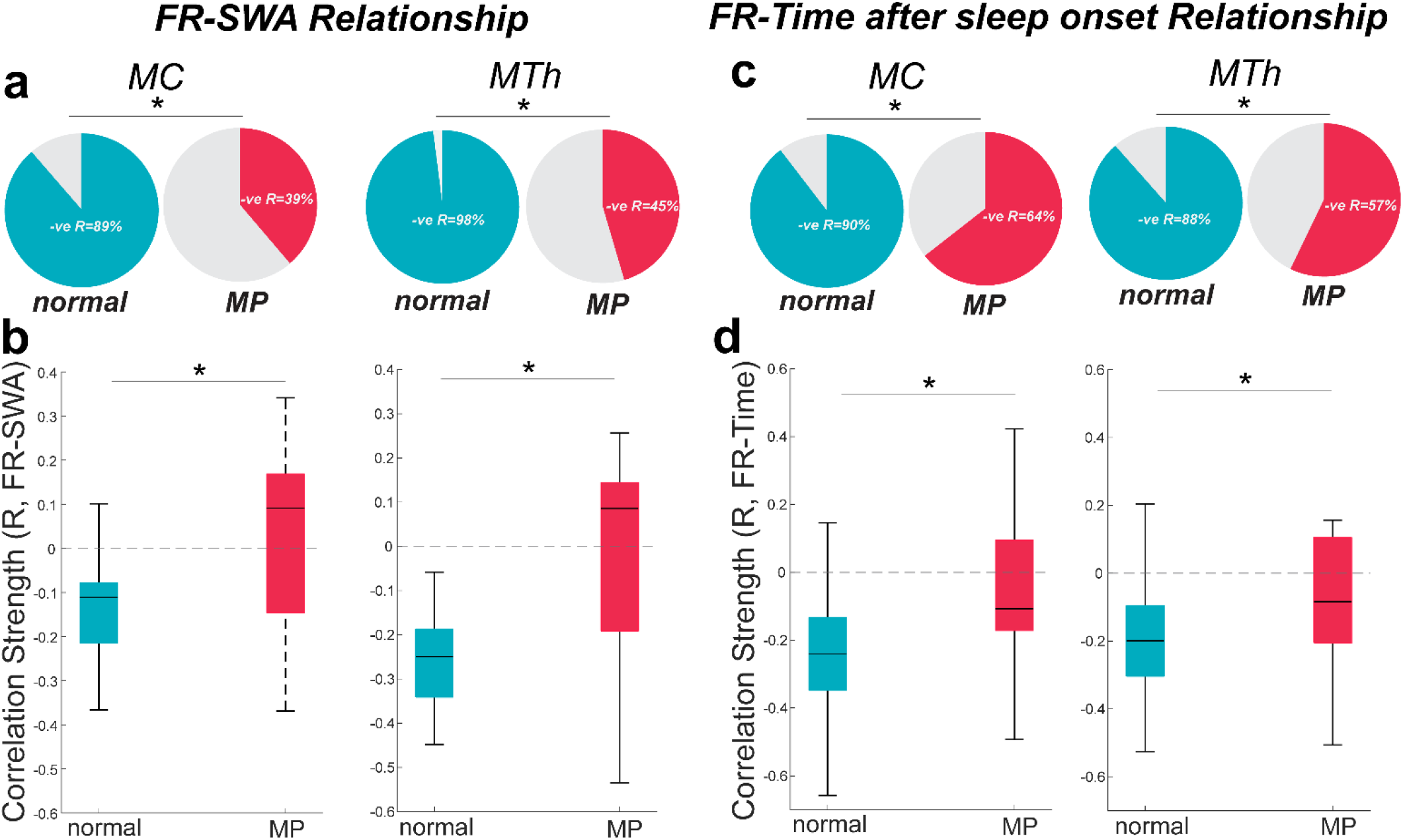
Out of all the MC and MTh neurons exhibiting significant firing rate (FR)-SWA correlation, the proportion of neurons showing significant negative FR-SWA correlation (R) was reduced in mild parkinsonism (MP) compared to normal for both MC (39/44 vs 19/49; normal vs MP) and MTh (49/50 vs 10/22; normal vs MP) neurons **(a)**. Moreover, the correlation polarity of MC and MTh neurons that exhibited significant FR-SWA correlation was switched to positive in MP as opposed to negative in normal **(b)**. Out of all the MC and MTh neurons exhibiting a significant correlation between FR and time (after the onset of sleep), the proportion of neurons exhibiting a negative FR-time correlation was reduced in MP compared to normal for both MC (43/48 vs 29/45; normal vs MP) and MTh (46/52 vs 8/14; normal vs MP) neurons **(c)**. Furthermore, the correlation strength of MC and MTh neurons that exhibited a significant correlation between FR and time after the onset of sleep differed between normal and MP conditions **(d)**.

The goal of this study was to improve our understanding of the neuronal mechanisms underlying sleep dysfunction associated with mild parkinsonism. Compared to normal, we found that in mild parkinsonism, neurons in the motor cortex and thalamus showed significant alterations in SWA amplitude, firing rate modulation from wake to sleep states, and progressive downscaling of firing rate during sleep. These data provide evidence that FRH, a characteristic of healthy sleep, is significantly disrupted in mild parkinsonism.

The synaptic homeostasis hypothesis^18,19^ predicts synaptic downscaling of neurons, evidenced by a decrease in neuronal firing from wake to sleep and a progressive decrease in firing rate from the onset of sleep (FRH). Our observations of reduction in neuronal firing of thalamic and cortical neurons from wake to sleep and progressive decrease from the time of sleep onset in the normal state align with this hypothesis. These neural mechanisms appear to be disrupted in mild parkinsonism as evidenced by our findings of marked attenuation in DM of thalamic and cortical neurons from wake to sleep as well as alterations in firing rate vs SWA and firing rate vs time correlations. Given the proposed importance of sleep FRH mechanisms in the facilitation of cognition and memory consolidation^3,20^, it might be hypothesized that dysfunctional sleep FRH plays a role in the difficulty faced by people with PD to acquire and retain new skills^21,22^. Future studies are required for objective characterization of the extent to which dysfunction in FRH during sleep is associated with a decline in learning and cognitive abilities.

A recent report suggests that impairment in the FRH mechanism could be associated with disruption in neuromodulator signaling in the basal forebrain^23^, which is critically involved in the regulation of sleep-wake states^24^. PD and chronic treatment of non-human primates (NHPs) with neurotoxin MPTP disrupt neuromodulator (e.g., dopaminergic, noradrenergic and cholinergic) signaling critical for sleep-wake regulation^25,26^. These studies and our findings taken together suggest that the dysfunction in FRH could be a common feature underlying disruption in neural circuits involved in sleep-wake regulation. Evidence of disrupted FRH in preclinical models of Alzheimer’s disease^8,27,28^ and Fragile X syndrome^29^ support the notion that disruption in FRH may underlie broad neurodegenerative disorders that impact the neural circuits involved in sleep-wake regulation and may not be specific to PD.

An alternative explanation for disrupted FRH noted in this study could stem from PD-related metabolic dysregulation in the thalamocortical network. Cortical metabolic activity, determined via PET imaging, is typically reduced during sleep compared to wake^30^, and impairment in wake-to-sleep modulation of cortical metabolic activity could underlie sleep dysfunction (e.g., insomnia)^31^. Cortical metabolic activity has been observed to be disrupted in both PD patients^32^ as well as in MPTP-treated NHPs^33^. The attenuation in modulation depth of neuronal firing from wake to sleep that we observed in PD could signify parkinsonism-related thalamocortical metabolic dysregulation during sleep^30,34^. This notion is supported by a recent study suggesting mitochondrial dysfunction, a major player in regulating metabolic activity, to be associated with disruption in FRH^35^. The relationship between metabolic dysfunction and FRH during sleep in PD has not been studied, and whether metabolic dysfunction plays a role in the disruption of FRH we observed in mild parkinsonism should be investigated by future studies.

We acknowledge our limited sample size as a study limitation. A distinct advantage of this study, however, is a within-subject comparison of neuronal activity during sleep-wake behavior before and after induction of parkinsonism that is not feasible in human subjects. Furthermore, NHP sleep architecture is analogous to that of humans^36^, and the induction of parkinsonism closely mimics sleep-wake dysfunction (sleep fragmentation, insomnia, excessive daytime sleepiness) commonly observed in people with PD^17^. Thus, the findings from NHP studies can have direct clinical implications and inform treatments for enhancing sleep quality in PD. In this study, we showed that, underlying sleep dysfunction associated with mild parkinsonism, the FRH of thalamic and cortical neurons is disrupted. Whether disruption in FRH mechanisms precedes sleep dysfunction associated with parkinsonism and if normalizing the FRH mechanism via therapeutics will prevent subsequent sleep dysfunction and impact disease progression remains an open question and warrants future investigation.

In summary, our study provided key insights into the mild parkinsonism associated alterations in the sleep FRH mechanism of thalamic and cortical neurons. These findings will be critical for the development of novel therapies^37^ to normalize FRH during the early disease stage, which can help improve sleep quality and mitigate the adverse effects of sleep dysfunction on the quality of life of people with PD.

## Methods

### Experimental Protocol and Data Acquisition

All procedures were approved by the University of Minnesota Institutional Animal Care and Use Committee and complied with the US Public Health Service policy on the humane care and use of laboratory animals. One aged adult female rhesus macaque NHP (23 years old) was used in this study. The subject was instrumented with a 96-channel microdrive (Gray matter research) with moveable microelectrodes targeting MTh and MC, enabling the sampling of different neurons across multiple nights. Simultaneous video and wireless neuronal recording of cortical and thalamic areas was obtained while the subject was in its home environment using a Tucker Davis Technology (TDT) neural data acquisition system fitted with wireless transceivers from Triangle Biosystems (TBSI) with a sampling rate of ∼24000 Hz. The subject was rendered mildly parkinsonian by administering four weekly low-dose (0.2-0.3 mg/Kg) intramuscular injections of the neurotoxin 1-methyl-4-phenyl-1,2,3,6-tetrahydropyridine (MPTP). The disease severity was determined using the Unified Parkinson’s Disease Rating Scale modified for NHPs (mUPDRS), which rates symptoms of bradykinesia, akinesia, rigidity, and tremor of the upper and lower limbs as well as food retrieval on the side contralateral to deep brain stimulation electrode implant. Each symptoms were rated on a scale of 0–3 (0 = naïve, 1 = mild, 2 = moderate, and 3 = severe), maximum score = 27^13^. A composite score of 3-9 was considered mild, 10-18 moderate, and 19-27 a severe parkinsonian state. The results presented in the manuscript are from 13 sessions of sleep recordings across naïve (n=6) and mild parkinsonian (n=7; mUPDRS: 5.75±0.51, mean±SD) conditions. The sleep recordings began at approximately 7 p.m. (lights off at 6 p.m.), and the first 2.5 hours of data, representing the first sleep cycle, were analyzed for evaluating FRH.

### Sleep staging and slow wave amplitude (SWA) quantification

Local field potentials from MC recording electrodes were sampled at 24kHz, bandpass filtered between 0.1 and 100 Hz, and further downsampled to 200 Hz. The processed LFP from MC electrodes were then averaged and normalized by z transformations. Slow wave amplitude was defined as the average amplitude calculated from the envelope of LFP filtered between 0.1 and 4 Hz computed using Hilbert transformation. The data was further divided into epochs of 5 seconds. Each 5-second epoch is first classified as whether belonging to a wake/sleep state based on video recording, movement artifacts, and slow wave amplitude. Quiescent wake periods were epochs during which the animal’s eyes were open, but minimal body movements were observed. Sleep periods were defined as periods devoid of movement and characterized by epochs exceeding 1 SD above the mean SWA in the quiescent wake period. Slow wave sleep (SWS) epochs were further identified as sleep epochs that exceeded 3 SD above the mean SWA in the quiescent wake period. The average SWA amplitude for each night was defined as the average SWA amplitude of all sleep epochs. All data processing was performed using custom scripts in MATLAB.

### Spike data extraction and quantitative analysis of firing patterns

Recordings from each electrode were filtered between (300–5000 Hz) and single units were isolated using PCA and visual inspection using Offline Sorter (Plexon Inc., Dallas, TX, USA). The firing rate of individual neurons was calculated across wake and sleep epochs. The average firing rate of a neuron in wake and SWS epochs was defined as the mean firing rate across all wake and SWS epochs, respectively. Depth of modulation (DM) was defined as the change in firing rate from wake to SWS. Pearson’s correlation between neuronal firing rate and time (since sleep onset) was calculated to understand whether the downscaling of neuronal firing as a function of time (since sleep onset) is preserved in PD.

### Statistical Analysis

The Wilcoxon rank sum (WRS) test was performed to evaluate differences in the DM and correlation strength of firing rate vs SWA amplitude and firing rate vs time since sleep onset in mild PD state compared to naïve. Fischer’s exact test was used to assess the proportion of neurons that reduced their firing rate from wake to sleep and the proportion of neurons that exhibited a negative firing rate-SWA and firing rate-time since sleep onset correlations. Pearson’s correlation coefficient was calculated to assess the relationship between firing rate vs SWA and firing rate vs time since sleep onset for individual neurons. Only the correlation strength for significant firing rate vs SWA and firing rate vs time correlation for MC and MTh neurons is presented in this study. Test results were considered significant at alpha=0.05.

## Acknowledgment

We would like to thank our animal core team of Claudia Hendrix, Hannah Baker, and Elizabeth McDuell as well as our veterinary and animal care colleagues at the University of Minnesota Research Animal Resources (RAR).

## Author Contribution

B.N., A.K.V., and L.A.J. conceived the research. Y.Y., E.M., and L.A.J. acquired data. B.N. analyzed data. B.N. and A.K.V. wrote the first draft of the manuscript. B.N. and A.K.V. performed statistical analysis. B.N. and A.K.V. made figures. B.N. and A.D. performed cortical and thalamic mapping. B.N., A.K.V., J.W., C.D.M., M.J.H., J.L.V., and L.A.J. interpreted results. J.L.V. and L.A.J. acquired funding and provided resources. All authors critically edited the manuscript and approved it for submission.

